# Dynamic but constrained: Repeated acquisitions of nutritional symbionts in bed bugs (Heteroptera: Cimicidae) from a narrow taxonomic pool

**DOI:** 10.1101/2025.06.13.659472

**Authors:** Václav Hypša, Jana Martinů, Sazzad Mahmood, Shruti Gupta, Eva Nováková, Ondřej Balvín

## Abstract

**Background:** Bed bugs (Heteroptera: Cimicidae), like other strictly blood-feeding insects, harbor obligate bacterial symbionts that supply essential nutrients (especially B vitamins) lacking in their blood diet. The primary obligate symbionts are transovarially transmitted *Wolbachia*, notable for having horizontally acquired biotin operon. In addition to these symbionts with the confirmed nutritional role, bacteria from the genera *Symbiopectobacterium* and *Tisiphia* have also been found to be transovarially transmitted in bed bugs. Their significance to the host fitness remains unclear; they are considered facultative, and thus likely non-essential. However, this understanding of bed bugs symbiosis is based exclusively on studies of the human-associated common bed bug (*Cimex lectularius*) and few related Cimicinae species. Virtually nothing is known about the diversity, origins, or metabolic roles of symbionts across the more than hundred described bed bug species.

**Results:** Using amplicon and metagenomic analyzes, we identified five different bacterial genera as potential bed bug symbionts: *Wolbachia*, *Symbiopectobacterium*, *Sodalis*, *Serratia*, and *Tisiphia*. The distribution of these bacteria among 13 bed bug species indicated at least 16 independent origins of symbiosis (some samples harboring multiple symbiont strains). A comparison of host and symbiont phylogenies suggested that some of these origins were followed by cospeciation. Not all bed bugs, however, harbored *Wolbachia*. In the subfamily Cacodminae, *Symbiopectobacterium* (previously known as facultative symbiont) was detected as the sole symbiont, suggesting its role as an essential, obligate symbiont. Analysis of 23 obtained genome drafts revealed considerable differences in their size and gene content, indicating that these bacteria have reached different stages in the evolution towards obligate symbiosis. As is common in symbiotic bacteria, the analyzed genomes have lost many biosynthetic capacities. A comparison of B-vitamin synthesis pathways showed that only two, riboflavin and lipoic acid, were preserved across all symbionts.

**Conclusions:** An overview of symbiosis across a broad phylogenetic span of bed bugs reveals that this insect family has undergone remarkably dynamic evolution, characterized by multiple independent symbiont acquisitions, episodes of cospeciations, and frequent co-occurrence of multiple symbionts within individual hosts. Interestingly, although these symbionts were acquired through multiple independent events, the majority belong to just three bacterial genera (*Wolbachia*, *Symbiopectobacterium*, and *Sodalis*), suggesting an unknown mechanisms underlying bed bug-symbiont specificity. Two aspects of this study warrant further investigation. First, the finding of *Symbiopectobacterium* as the sole obligate symbiont in Cacodminae suggests that expanding the taxonomic sampling may reveal an even more complex structure of symbiosis within bed bugs. Second, the inconsistencies and uncertainties in evaluating the functionality of the biotin synthesis indicate that further research will be necessary to better understand the evolution of this B vitamin pathway in symbiotic bacteria.

## Introduction

Many insect groups maintain obligate association with symbiotic bacteria that supplement essential nutrients missing from their diets (Douglas 2015). While some of these associations exhibit remarkable long-term stability (e.g., the aphid-*Buchnera* relationship which has persisted over a hundred million years of cospeciation; Moran and Baumann 2000), others undergo a dynamic process of symbiont acquisitions, losses, and replacements (Sudakaran, Kost and Kaltenpoth 2017). In insects that feed exclusively on vertebrate blood, the presence of obligate symbionts is critical for host viability and fitness, as these symbionts provide B vitamins that are deficient in the blood diet (Duron and Gottlieb 2020). Studies across various haematophagous insects have shown that these obligate mutualists originated from a diverse array of bacterial taxa (Rio, Attardo and Weiss 2016, Duron and Gottlieb 2020).

One such exclusively hematophagous group is the bed bugs, the heteropteran family Cimicidae, which comprises over a hundred described species, primarily associated with bats and birds (Reinhardt and Siva-Jothy 2007). However, for practical reasons, the most extensively studied cimicid species is the human-associated common bed bug (*Cimex lectulari*us). Three bacterial groups have so far been demonstrated or suggested to be transovarially transmitted symbionts in this species. The most prominent is a strain of the widespread insect symbiont *Wolbachia*. This bacterium, best known for manipulating insect reproduction (Werren, Baldo and Clark 2008), has been first identified in two cimicid species by Rasgon and Scott (2004), and its role of an obligate symbiont in *C. lectularius* was subsequently documented by several studies (Hosokawa et al. 2010, Nikoh et al. 2014, Hickin, Kakumanu and Schal 2022, Poulain et al. 2024). The other two symbionts are considered facultative and likely non-essential. The first is a bacterium related to the symbiont of *Euscelidius variegatus*, initially described from *Cimex lectularius* (Hypsa and Aksoy 1997), and later shown to belong to the genus *Symbiopectobacterium* (Nadal-Jimenez et al. 2022). The second is a *Rickettsia*-related *Candidatus* Tisiphia (Thongprem et al. 2020, Cagatay et al. 2025). The significance of these two facultative symbionts for the host fitness remains unclear.

Several lines of evidence strongly support the view that *Wolbachia* are obligate symbionts essential for bed bug development and reproduction: their ubiquity across bed bug populations, efficient transovarial transmission (Hosokawa et al. 2010, Cagatay et al. 2025), and presence of a horizontally acquired biotin operon (Nikoh et al. 2014). A typical consequence of such an intimate host-symbiont association is a shared coevolutionary history, reflected in congruent host and symbiont phylogenies. A screening study by Balvin et al. (2018) indeed revealed a partial coevolutionary signal between *Wolbachia* and their Cimicinae hosts. However, this cospeciation pattern did not include all Cimicidae associated *Wolbachia*, suggesting that the symbiont evolution within this host group is more complex than a model of strict vertical transmission and coevolution. This finding aligns with earlier results by Sakamoto, Feinstein and Rasgon (2006) who detected *Wolbachia* strains from two different supergroups, F and A, in cimicid museum specimens. Furthermore, in *C. lectularius*, genome-wide analysis has revealed extensive lateral gene transfers from several bacterial genera, including known insect symbionts *Arsenophonus*, *Sodalis*, and *Hamiltonella* (Benoit et al. 2016). These genetic remnants may represent relics of ancient, now-lost symbiotic association, as suggested in other insect systems (Husnik and McCutcheon 2016). Indirect support for this hypothesis of dynamic evolution in cimicid-symbiont associations comes from comparisons with other obligate haematophages, which harbor diverse microbiomes of complex evolutionary histories (Ríhová et al. 2021, Sochova et al. 2017).

To directly test the hypothesis that symbioses in Cimicidae have followed a complex and dynamic evolutionary trajectory characterized by multiple independent origins across diverse bacterial genera, we analyzed a broad set of cimicid specimens using both amplicon and metagenomic sequencing. Amplicon sequencing served primarily as a screening tool to assess the distribution of symbiotic taxa across the Cimicidae phylogeny. Metagenomic sequencing was used to explore the genomic characteristics and metabolic capacities of the symbionts, focusing on 14 samples representing 13 species from five subfamilies: Afrocimicinae, Haematosiphoninae, Cacodminae, Cimicinae, and the basal Primicimicinae.

## Results and discussion

### Amplicon screening

Amplicon screening revealed several dominant OTUs belonging to well-known symbiotic taxa (Figure 1). The most abundant taxon, found in 10 out of 13 species, was assigned to the genus *Wolbachia* represented by three OTUs in our dataset. The nucleotide identity of the best blast hits suggests that these OTUs likely correspond to different *Wolbachia* symbionts from three supergroups (*Wolbachia* F, E, and B; Supplementary table S1). The distribution pattern of these OTUs (Figure 1) further suggest dual *Wolbachia* infection for at least two host species, *Cimex hirundinis* and *Cyanolicimex patagonicus.* The genus *Symbiopectobacterium* is represented by a single OTU dominating microbiomes of *Cacodmus* spp. and *Leptocimex duplicatus*. This *Symbiopectobacterium* OTU is also detected alongside with *Wolbachia* in *Cimex hemipterus* and *Cimex lectularius* specimens collected from human hosts. In contrast, the distribution of *Serratia* and *Tisiphia* (torix Rickettsia) symbionts is highly host-specific, each being restricted to a single species in our dataset, i.e., *Paracimex* sp. and *Afrocimex constrictus*, respectively. The last symbiotic taxon identified, belonging to the genus *Sodalis*, is represented by 4 OTUs and exhibits the most complex distribution pattern. Two *Sodalis* OTUs associated with *Cyanolicimex patagonicus* display surprisingly low nucleotide identity (92,75%). In *Psitticimex uritui* microbiome, the other two *Sodalis* OTUs vary among the screened individuals. The presence of multiple OTUs for the same symbiotic taxon can, in principle, reflect either a true biological pattern or a methodological artefact (e.g., sequencing errors). However, because our amplicon data were processed using a highly conservative pipeline that retains only the highest-quality reads (see Materials and Methods), a biological explanation is more likely. The observed diversity may thus result from coinfection by distinct symbiont lineages or from intragenomic heterogeneity of 16S rRNA gene copies, especially in *Sodalis* symbionts known to carry up to seven rRNA operons (Oakeson et al. 2014). Since nucleotide identities among OTUs for both *Sodalis* and *Wolbachia* symbionts range from 92.75% to 96.75%, we propose that these OTUs represent distinct symbiotic lineages.

**Figure 1.**
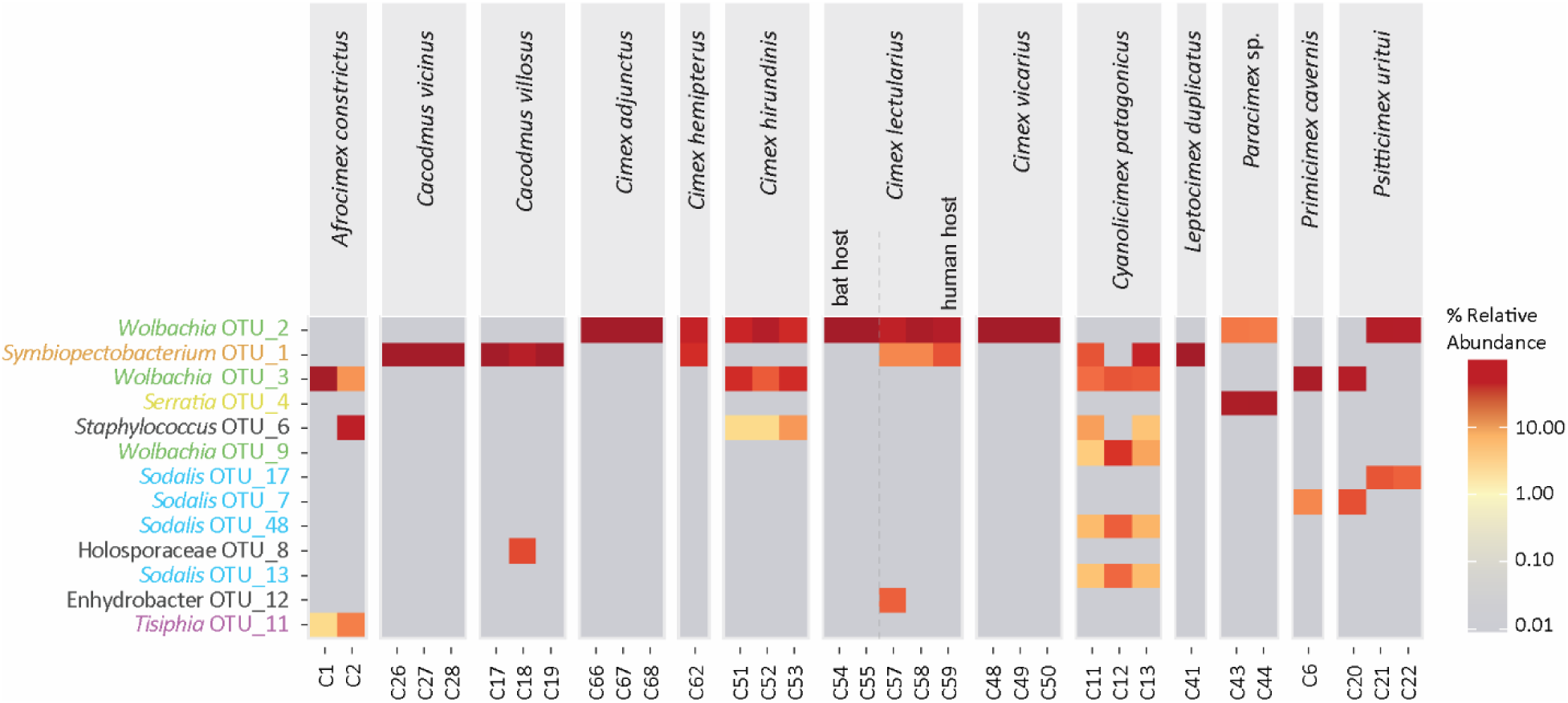
Distribution of first 13 most abundant OTUs found across 33 cimicid samples. Symbiotic taxa are color-coded to highlight their (co-)occurrence and relative abundance across the dataset.

### Metagenomic assemblies and the drafts of symbionts’ genomes

The Illumina short-reads sequencing data obtained for 14 Cimicidae samples (13 species), along with the main characteristics of their metagenomic assemblies, are summarized in Supplementary table S2. Screening these assemblies using bacterial gene queries led to identification of 23 genome drafts corresponding to known symbiotic taxa, as indicated by the amplicon screening (Table 1). Among these, 14 *Wolbachia* strains were detected in 11 host samples, with three samples exhibiting double infections (C51-*Cimex hirundinis*, C12-*Cyanolicimex patagonicus,* and C21-*Psitticimex uritui*). The majority of these genomes ranged in size approximately 1 to 1.5 Mbp, aligning well with the sizes reported for complete *Wolbachia* genomes available in the NCBI GenBank. In addition to *Wolbachia*, genomes of *Symbiopectobacterium, Sodalis, and Serratia* were identified in four, three, and one assembly, respectively. In C1-*Afrocimex constrictus*, we also detected fragments of *Thisiphia* genome. However, these sequences had very low coverage (1.7 to 8.6) and BUSCO completeness (Supplementary table S3) and were not included in the subsequent analyses. The *Symbiopectobacterium* and *Sodalis* genomes were larger than those of *Wolbachia*, ranging approximately between 2 and 3 Mbp. Assembly quality varied substantially among samples, from a complete, closed genome (*Wolbachia* C44-*Paracimex* sp.) to draft assemblies comprising hundreds of scaffolds. Genome completeness, assessed by BUSCO using ricketssia_db12 database, was high for most of the *Wolbachia* genomes, exceeding 90% in all but two samples (Supplementary table S3). For *Sodalis* and *Symbiopectobacterium*, completeness values fell below 50% in some samples when assessed with pectobacteriaceae_db12 database but increased substantially (exceeding 80%) when evaluated with the enterobacteriales_db12 and bacteria_db12 reference datasets. The GC content varied greatly among the genomes, ranging from 30% to 60%. Genome size and GC content were largely correlated, with a distinct separation between *Wolbachia* (lowest GC and shortest genomes) and the other symbionts.

**Table 1.**
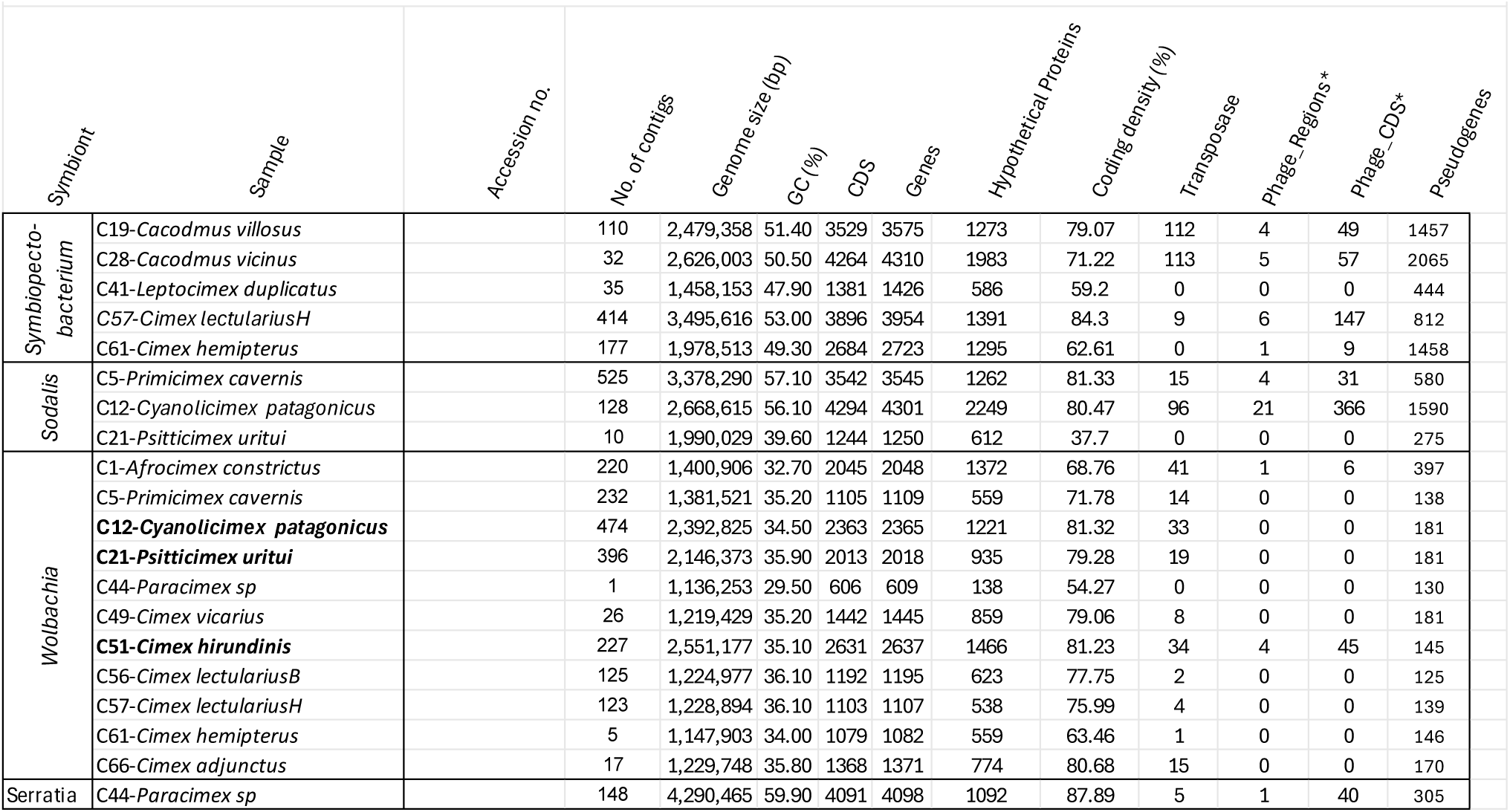
Main characteristics of the symbionts’ genomes. Genomes printed in bold stands for double infection with two different strains (see methods for details). * = predicted by Phastest https://phast.

In addition to standard bacterial coding sequences (CDSs), the genomes exhibited a diverse array of mobile genetic elements (MGEs), notably transposases and phage-related sequences. An intriguing pattern was observed among the five genomes of *Symbiopectobacterium*. Two strains (C28-*Cacodmus vicinus* and C19-*Cacodmus villosus*) exhibited the highest numbers of transposases. A comparison with the large-genome *Symbiopectobacterium purcellii* (strain SyEd1, NZ_CP081864.1) suggests that genome reduction in these two strains is primarily due to the loss of entire genomic regions, often adjacent to a transposase elements (Supplementary figure S1). Given the limited number of *Symbiopectobacterium* genomes available for the comparison, any interpretation remains speculative; however, a plausible evolutionary scenario can be proposed. The largest genome (C57-*C. lectularius*) harbors relatively few transposases, potentially representing an early stage of symbiont adaptation. Subsequent stages appear to involve acquisition of numerous transposases, facilitating the deletion of non-essential genomic regions, as evidenced by the intermediate genomes (C28-*Cacodmus vicinus* and C19-*Cacodmus villosus*). This phase of rapid genome degeneration is also characterized by an elevated number of pseudogenes. In the advanced stage, genomes not only become more compact but also exhibit a significant reduction in transposase genes, as observed in the two smallest genomes (C61-*C. hemipterus*, C41-*Leptocimex duplicatus*; Supplementary figure S1).

### Phylogenetic relationships and coevolutionary patterns

#### Cimicidae phylogeny

Both maximum likelihood (ML) and Bayesian inference (BI) phylogenetic analyses, based on an alignment of 35,995 amino acid residues derived from nuclear-encoded proteins, yielded identical topologies with strong supports (all posterior probabilities of 1.0 and bootstrap values 100; Figure 2). This topology differs in two key aspects from the phylogeny published by Roth et al. (2019), which was based on four genes. First, Cacodminae and Haematosiphoninae form a paraphyletic group relative to the monophyletic Cimicinae. Second, within Cimicinae, *Paracimex* branching within *Cimex*, alternating the internal topology of the subfamily. Notably, this revised topology is not only well supported, but also congruent with the phylogeny of the corresponding *Wolbachia* strains (highlighted in green; compare to Figure 3).

**Figure 2.**
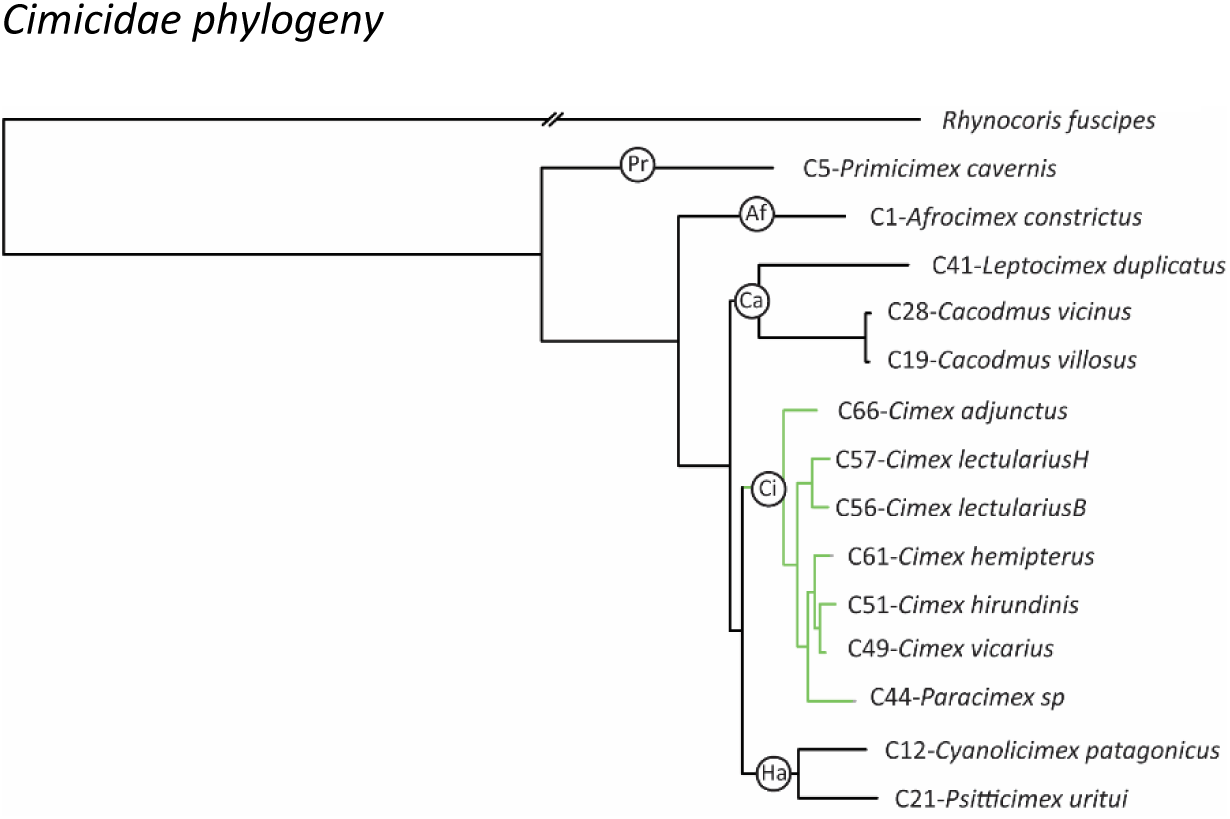
Phylogenetic relationships among Cimicidae samples inferred by ML from an alignment of 35,995 amino acid residues derived from nuclear-encoded proteins. All bootstrap support values were 100. BI recovered an identical topology, with all posterior equal to 1.0. Af=Afrocimicinae, Ca=Cacodminae, Ci=Cimicinae, Ha=Haematosiphoninae, Pr=Primicimicinae. Green branches indicate the Cimicinae-Wolbachia cospeciation.

**Figure 3.**
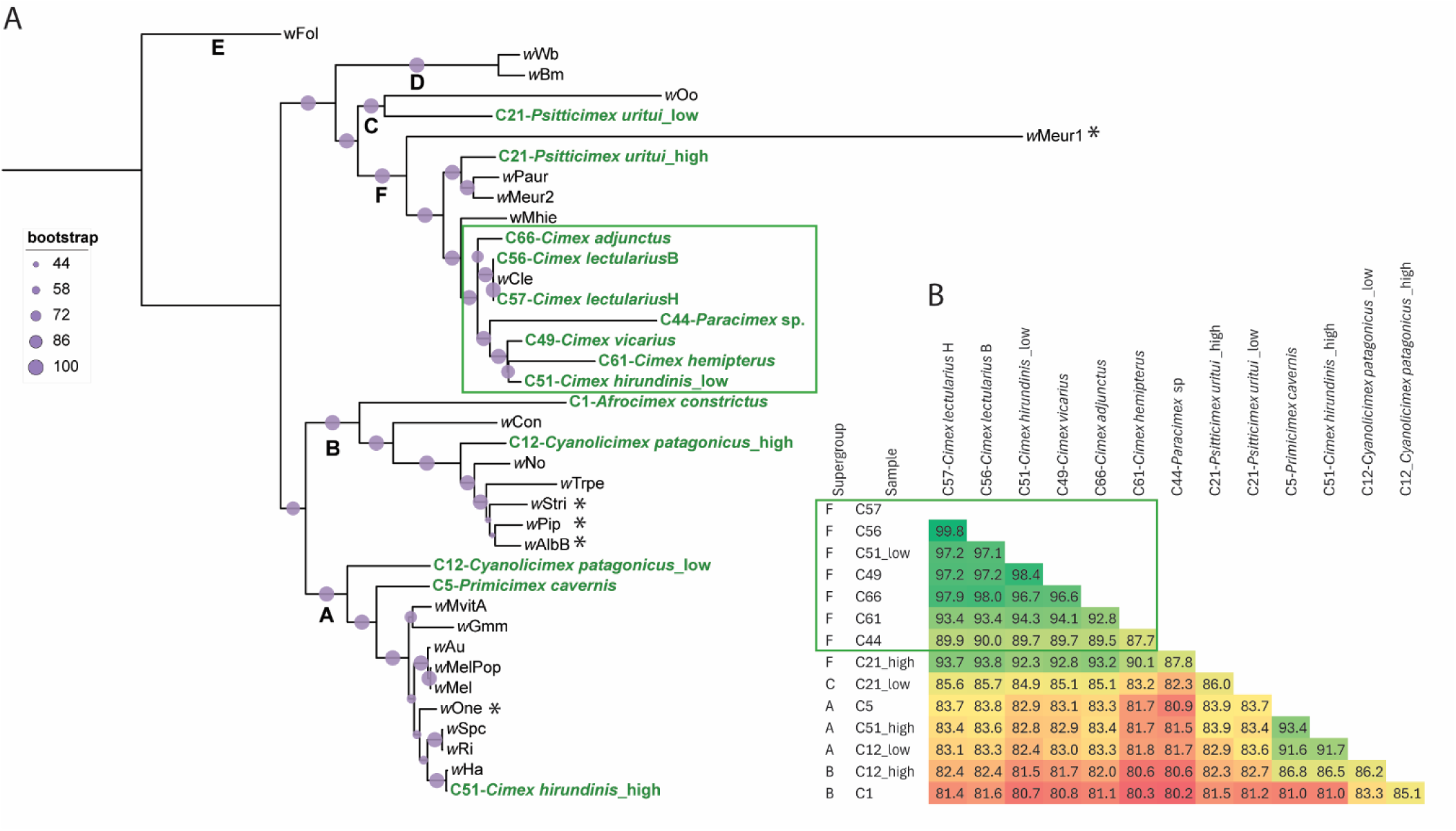
Phylogenetic relationships and genomic similarities among *Wolbachia* strains. A) ML phylogenetic tree inferred from from concatebnated matrix of 115 single-copy orthologs (27,850 amino acid residues). Letters at the nodes indicate *Wolbachia* supergroups. Strains identified in this study from cimicid species are highlighted in bold green. The designations “high” and “low” indicate differences in sequencing coverage between co-occurring strains in the same host. The green rectangle highlights the coevolving lineage. Asterisks mark branches whose positions differ in BI analysis. B) Average nucleotide identity (ANI) of *Wolbachia* strains associated with cimicid hosts. The green rectangle highlights the coevolving lineage. Accession numbers for all included taxa are provided in Supplementary table S4.

#### Symbionts phylogenies

##### Wolbachia

Phylogenetic analyses using ML and BI based on 115 single-copy orthologs (27,850 amino acid residues), recovered largely identical topologies, with only few minor differences (Figure 3; Supplementary figure S2). These analyses revealed substantial phylogenetic diversity among the *Wolbachia* strains from cimicid hosts. Strains belonging to supergroup F, associated with Cimicinae species (including *Paracimex*), formed a well supported monophyletic cluster whose topology mirrored that of their host, as recovered in this study (Figure 2), consistent with a previously proposed coevolutionary scenario (Balvin et al. 2018). In contrast, the remaining seven strains, affiliated with other supergroups, clustered with strains from unrelated insect hosts and showed no correspondence with cimicid phylogeny. In four samples exhibiting double infections, the co-occurring *Wolbachia* strains occupied distinct positions in the phylogenetic tree, indicating divergent evolutionary origins. Genome-wide average nucleotide identity (ANI) analyses supported these patterns: within the coevolving supergroup F cluster, ANI values exceeded 90% (with the exception of *Paracimex*-associated strain), whereas ANI values between co-occurring strains from the same host ranged from 82% to 86%, confirming their distinct phylogenetic affiliations.

##### Symbiopectobacterium

Both ML and BI analyses of five *Symbiopectobacterium* strains, based on 340 single-copy orthologs (totaling 90,300 amino acid residues), recovered an identical topology that clustered all five cimicid-associated strains together with a nematode-associated strain (*Candidatus* Symbiopectobacterium sp. ‘North America’) (Figure 4). Despite their apparent close phylogenetic relationships, the positions of the cimicid-associated strains suggest at least four independent acquisitions of *Symbiopectobacterium* by bedbugs. Only in one instance (a pair of sister *Symbiopectobacterium* strains C28 *C. vicinus* and C19 *C. villosus*) did the symbiont phylogeny correspond with the host relationship, as both were isolated from closely related Cacodmus species. The remaining three strains clustered without any apparent correspondence to their hosts’ phylogeny. Although both ML and BI analyses recovered the same topology with well supported nodes, it is noteworthy that at least two strains (C41-*Leptocimex duplicatus* and 61-*Cimex hemipterus*) formed long branches, which could potentially result in topological artifacts.

**Figure 4.**
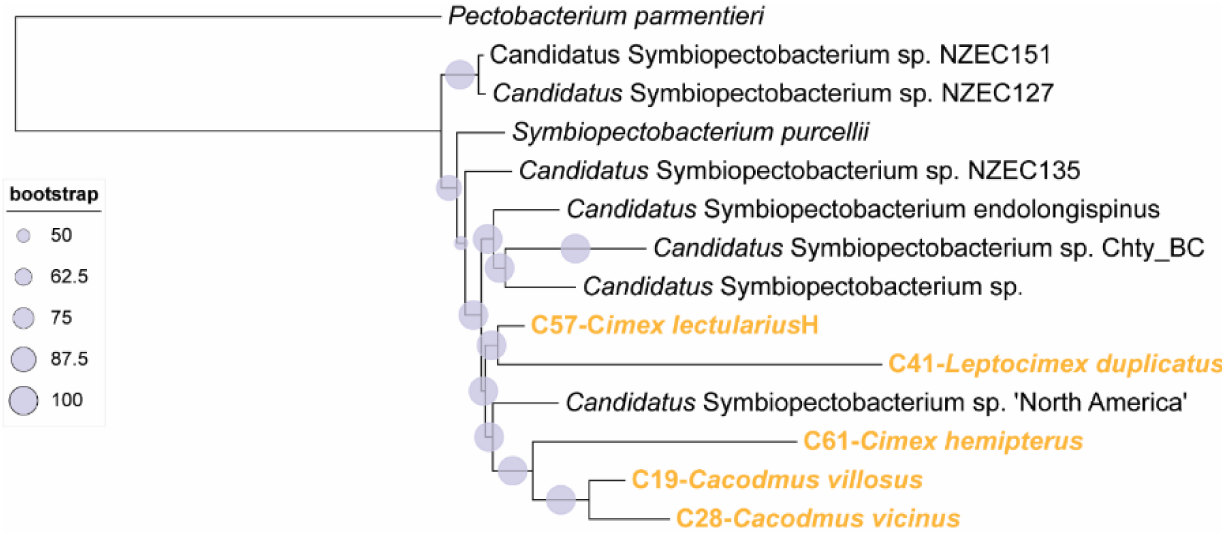
Phylogenetic relationships of the Symbiopectobacterium strains inferred by ML analysis of 340 single-copy orthologs (totalling 90,300 amino acid residues). Strains identified in this study from cimicid species are highlighted in bold orange. The BI analysis recovered an identical topology, with all posterior probabilities equal to 1. Accession numbers for all included taxa are provided in Supplementary table S4.

##### Sodalis

The phylogenies inferred by ML and BI analyses based on 41 single-copy orthologs (11,296 amino acids) differed in several aspects (Figure 5) but both consistently placed the bedbug-associated *Sodalis* strains into two distinct clusters: first comprising C12-*Cyanolicimex patagonicus* and C21-*Psitticimex uritui*, and the second C5-*Primicimex cavernis*. This well-supported topology suggests at least two independent origins of the Cimicidae-*Sodalis* symbiosis. Furthermore, the phylogenetic affiliations and branch lengths within these two clusters indicated different characters of the symbiotic associations. The C12-*Cyanolicimex patagonicus* and C21-*Psitticimex uritui* strains were related to other insect symbionts and possessed relatively long branches (particularly the latter one), in contrast to C5-*Primicimex cavernis*, which was placed on very short branch among the strains of *S. praecaptivus*. Evolutionary interpretation of the Cimicidae-*Sodalis* association is, however, complicated by an inconsistent position of the two long-branched strains associated with the Haematosiphoninae species. While the ML analysis positioned them as sister taxa, BI tree placed them in a paraphyletic arrangement, where C21-*Psitticimex uritui* strain was related to another particularly long-branch bacterium, the *Sodalis* CWE strain described from chewing louse (Alickovic, Johnson and Boyd 2021). Both these alternative placements were only supported by short branches and low bootstrap and posterior probability values, indicating potential instability in these relationships.

**Figure 5.**
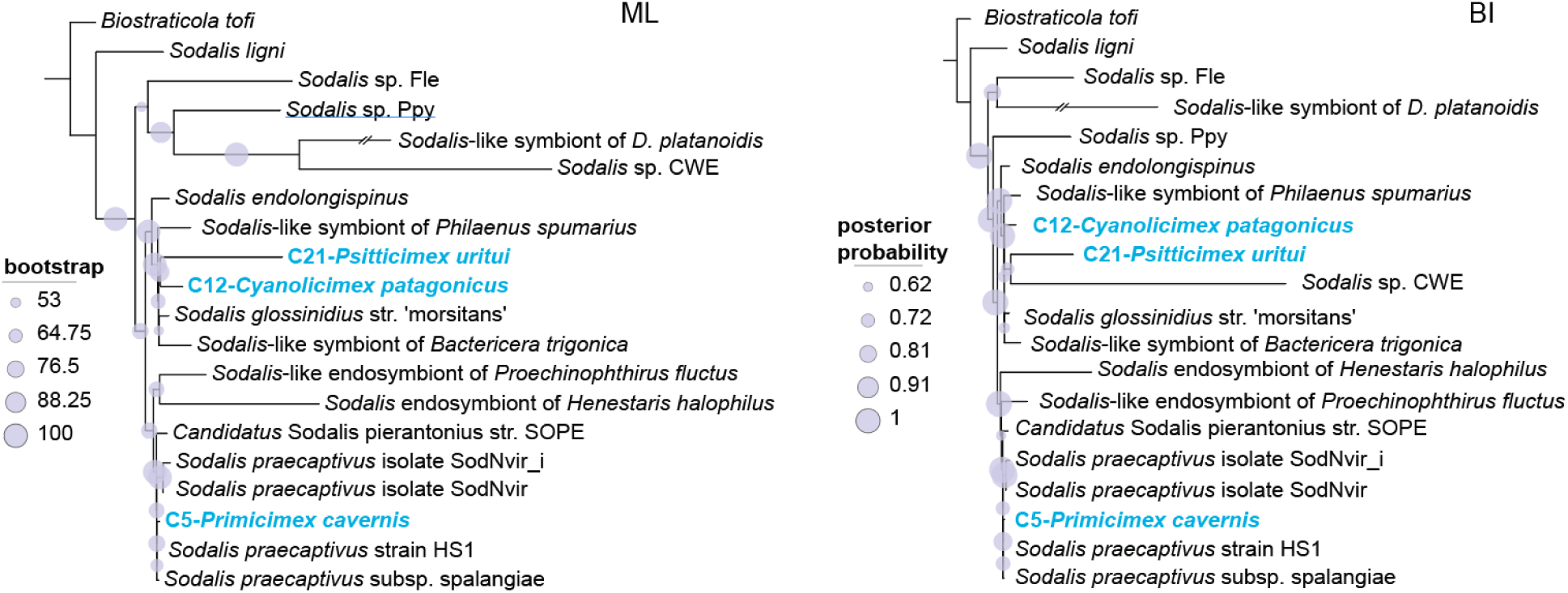
Phylogenetic relationships of the *Sodalis* strains inferred by ML and BI analyses of 41 single-copy orthologs (11,296 amino acids). Accession numbers for all included taxa are provided in Supplementary table S4.

##### Serratia

Both analyses based on 1,271 single-copy orthologs (394,084 amino acids), consistently placed the sole identified *Serratia* strain (C44-*Paracimex* sp.) in a clade with the nematode symbiont *Serratia nematodiphila*, a nematode-associated bacterium, and *Serratia* sp. strain BNK-11 of an unclear origin (Figure 6). This grouping was distinct from the clade containing the aphid symbiont *S. symbiotica*.

**Figure 6.**
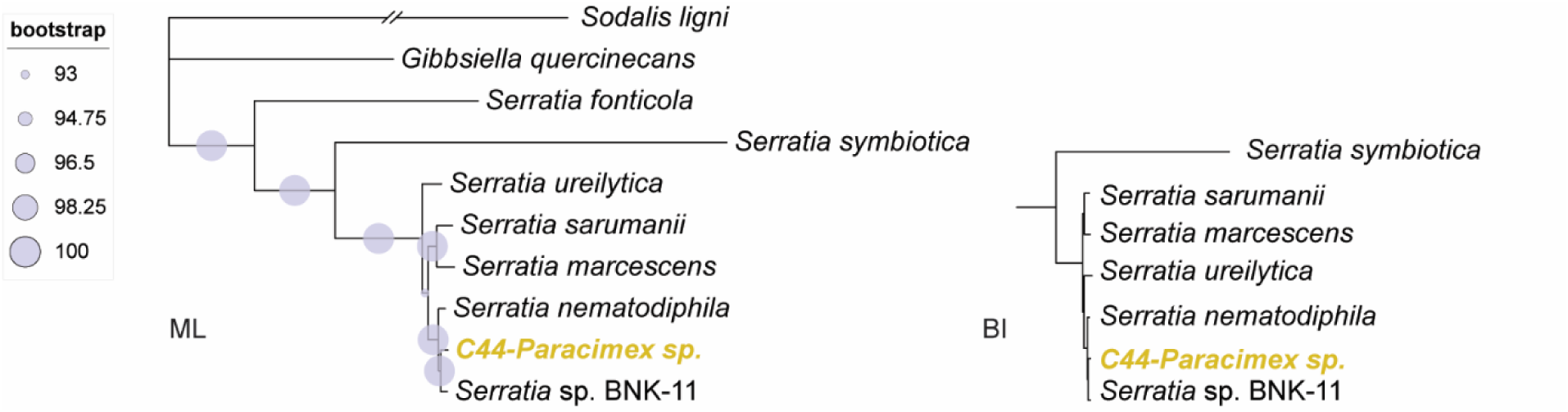
Phylogenetic placement of the *Serratia* symbiont associated with *Paracimex* sp. Based on 1,271 single-copy orthologs (394,084 amino acids), both ML and BI analyses inferred identical topologies, with only minor difference observed within the cluster of the short-branched Serratia strains. Accession numbers for all included taxa are provided in Supplementary table S4.

#### Metabolic capacity

Comparison of metabolic capacities across the analyzed symbionts (overview in Supplementary table S5) reveals a pattern shaped by two main factors. First, in many metabolic pathways, the presence or absence of genes is influenced by the symbiont’s phylogenetic position. This is particularly evident when comparing the two phylogenetically distant groups, *Wolbachia* (Alphaproteobacteria) and *Sodalis/Symbiopectobacterium* (Gammaproteobacteria). Notable examples include several amino acid biosynthesis pathways, ABC transporters, two component systems, and lipopolysaccharide synthesis. Second, metabolic capacity is determined by the extent of genome degradation. This is clearly illustrated by the *Symbiopectobacterium* strains C41-*Leptocimex duplicatus* and C61-*Cimex hemipterus* which exhibit considerably less complete metabolic pathways compared to the other *Symbiopectobacterium* strains (reflected also in smaller size of the two genomes). This reduction appears to reflect genuine genomic degradation rather than incomplete assemblies, as evidenced by lower GC content and long branches these strains form in phylogenetic trees, both indicative of ongoing genomic degeneration.

A detailed assessment of B vitamins synthesis (Figure 7) revealed that only two pathways, riboflavin and lipoic acid, were universally functional in all symbionts, including different symbiont genera inhabiting a single host. The significance of riboflavin pathway deduced from its functionality in all samples agrees with some previous studies. In comparative genomic study of *Wolbachia* strains, synthesis pathway for riboflavin was the only synthetic capability of B vitamins conserved among diverse insects, and its importance for bed bug growth, survival, and reproduction was demonstrated experimentally (Moriyama et al. 2015). Likewise, the riboflavin synthesis has been universally functional in all 11 analyzed strains of another symbiotic bacterium, the genus *Arsenophonus* associated with hippoboscids (Martin Říhová et al. 2023). Among the 22 symbionts analyzed in our study (with *Wolbachia* double infections treated as single strain for metabolic purposes), 13 lacked the gene annotated for the dephosphorylation step in the standard riboflavin pathway, as defined in KEGG database. However, this step can be catalyzed by several alternative enzymes, and similar “gaps” have been reported in other symbiotic bacteria (Rihova, Vodicka and Hypsa 2025). Given the retention of all other genes in the pathway, this suggests that riboflavin synthesis remains functional, despite the missing canonical dephosphorylation gene. A third pathway, folate biosynthesis, was found to be functional in most symbionts. The two exceptions were the *Wolbachia* strains from *Paracimex* sp. and *C. adjunctus*. In the case of *Paracimex* sp., the co-occurring *Serratia* symbiont retains all functional B vitamins pathways, leaving the *C. adjunctus* sample the only one without apparent capacity for folate synthesis.

**Figure 7.**
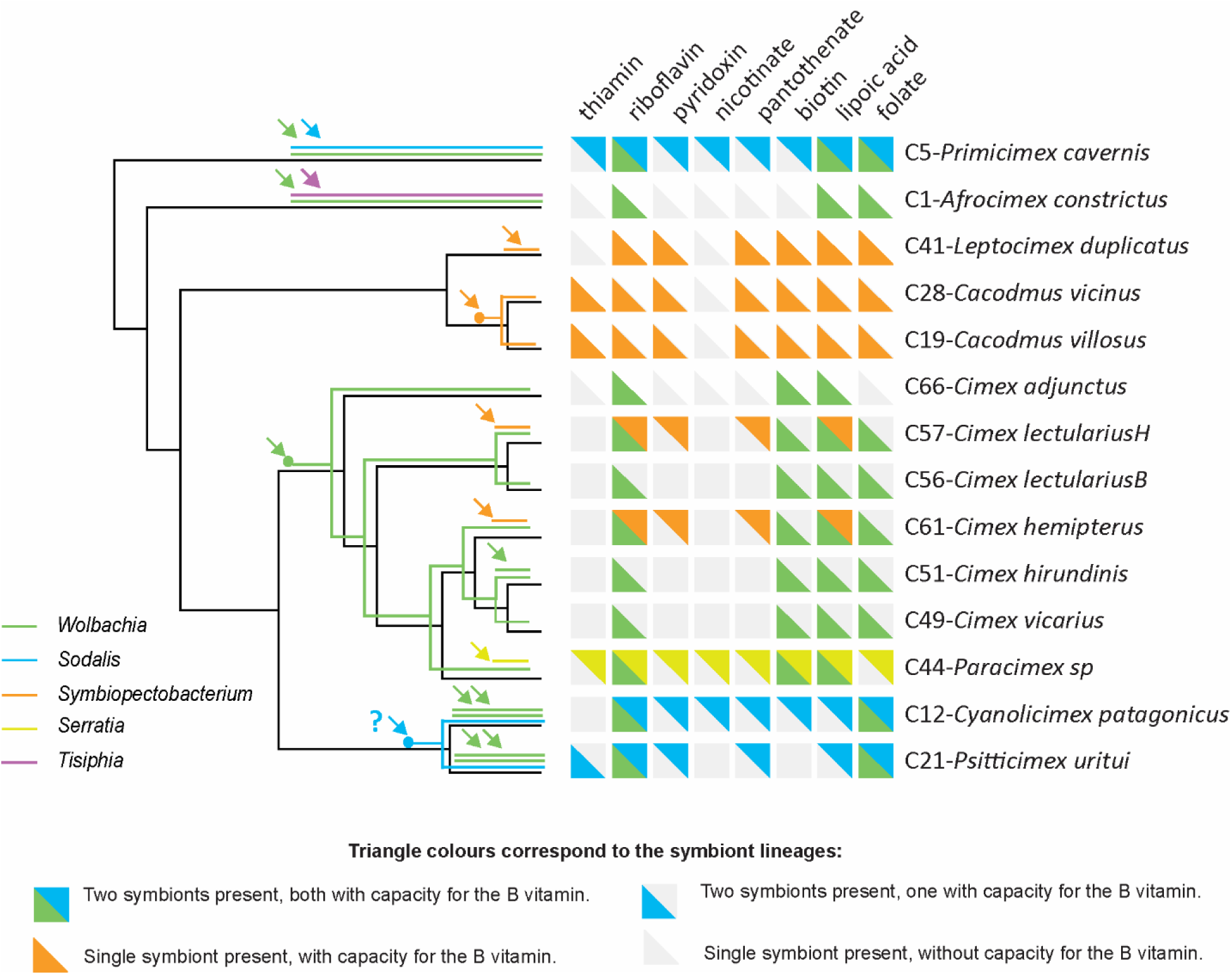
Symbiont origins and metabolic capacities for B vitamins mapped on phylogenetic relationships of the hosts. Arrows point to hypotethical independent origins. Dots indicate hypothetical cospeciation of the host and the symbiont (the question mark in Sodalis lineage points out uncertainty in respect to cospeciation versus two independent origins).

Three B-vitamin biosynthesis pathways (thiamine, pyridoxal, and pantothenate) were non-functional in all *Wolbachia* strains but were largely retained in the other symbionts. In particular, pyridoxal and pantothenate pathways seem to be functional in all strains of *Symbiopectobacterium*, *Sodalis,* and *Serratia*. In the C57-*C. lectularius* sample, thiamine synthesis may be potentially functional if the partial biosynthetic capacities of the two co-resident symbionts, *Wolbachia* and *Symbiopectobacterium*, are combined. In contrast, the capacity for nicotinate and cobalamine biosynthesis were absent in most or all symbionts, respectively. The most intriguing pattern was observed in biotin synthesis. Biotin is commonly considered essential for obligate blood-feeding insects and supplemented by the symbionts (Duron and Gottlieb 2020). This seems to be supported by independent acquisitions of the biotin operon by horizontal gene transfer (HGT) in several symbiotic bacteria (Nikoh et al. 2014, Rihova et al. 2017). Furthermore, experimental study demonstrated that supplementation with biotin and riboflavin effectively restored the fitness of bed bugs following the elimination of their symbionts (Moriyama et al. 2015). In our data, the biotin pathway was functional in most, but not all, of the analyzed symbionts. One line of evidence highlighting the potential importance of biotin supplementation comes from its distribution in the *Symbiopectobacterium* strains. The pathway was present in strains from all three Cacodminae species, where *Symbiopectobacterium* is the sole symbiont, but absent in both strains from Cimicinae species, where biotin is instead produced by co-occurring *Wolbachia*. Moreover, the genome of the *Wolbachia* strain C44-*Paracimex* carries duplicated biotin operon, with the two copies complementing each other: in one copy, the bioB gene is disrupted by a premature stop codon, while the other copy contains functional bioB and disrupted BioF, bioD, and bioA (Supplementary figure S3). The signs of gene degradation within this pathway in some other *Wolbachia* strains further complicate the interpretation on the essentiality of biotin. Although these strains retain the biotin operon (originally obtained by HGT) several genes appear truncated or fragmented due to premature stop codons, and were not annotated as functional by BlastKoala. In at least two samples (C1-*Afrocimex constrictus* and C21-*Psitticimex uritui*), the biotin pathways of their symbionts are very likely non-functional.

#### Symbionts’ origins and coevolutionary history

An overview of the 14 Cimicidae samples reveals that their associations with symbiotic bacteria have undergone remarkably dynamic evolution, with multiple independent symbiont acquisitions and frequent co-occurrence of multiple symbionts within individual hosts (Figure 7). Interestingly, *Symbiopectobacterium*, previously known as facultative symbiont co-occuring with obligate *Wolbachia*, was detected in this study as the sole symbiont in three Cacodminae species, entirely lacking *Wolbachia*. Mapping the symbiont associations onto the cimicid phylogeny suggests at least 16 independent origins (Figure 7). Only three of these origins show evidence of subsequent coevolution across host species boundaries: seven strains of *Wolbachia* associated with Cimicinae, two *Symbiopectobacterium* strains from the two *Cacodmus* species, and two *Sodalis* strains from the Haematosiphoninae species. The first two cases are straightforward: the symbionts form well-supported monophyletic lineages, each associated with a group of closely related hosts. The pattern observed in *Sodalis*, however, is less clear. The two host genera (*Cyanolimex* and *Psitticimex*) belong to the same subfamily but are not sister taxa, or particularly closely related genera. If the observed pattern does reflect a process of host-symbiont cospeciation, the full coevolutionary history of this *Sodalis* lineage likely involves additional genera/species within the Haematosiphoninae and their associated *Sodalis* strains. This interpretation is also consistent with the long branches of both coevolving *Sodalis* strains in our phylogenetic analysis (C12-*Cyanolicimex patagonicus* and C21-*Psitticimex uritui*).

Considering the high number of symbiont acquisitions, it is noteworthy that they involve only a limited set of bacterial taxa. The majority are represented by different strains of *Wolbachia*, with eight independent origins, followed by four origins from *Symbiopectobacterium*, two from *Sodalis*, and one from *Serratia* and *Tisiphia*. The distribution, coevolutionary patterns, and genome characteristics of these bacteria suggest that the different strains have reached various stages in their symbiogenesis. These stages are reflected in traits such as genome reduction, metabolic degradation, shift of nucleotide composition, and presence of mobile elements.

The observed patterns also raise several important questions for future research. One key direction is to better understand the overall diversity of symbionts across Cimicidae, which will require addressing phylogenetic gaps in the host sampling. The importance of this step is underscored by the identification of *Symbiopectobacterium* as the sole symbiont in Cacodminae species, extending the known obligate symbionts in Cimicidae to at least two genera, *Wolbachia* and *Symbiopectobacterium*. The dataset analyzed in this study covers all currently recognized subfamilies of bed bugs, except for Latrocimicinae (a group consisting of a single bat related species). Within the best-represented subfamily, Cimicinae (with six included species), our sample allowed for demonstration of two patterns: (i) *Wolbachia*-Cimicinae cospeciation across six host species, and (ii) several independent origins of *Wolbachia, Symbiopectobacterium*, and *Sodalis* strains. However, a more comprehensive sampling will be essential to gain deeper insight into the symbiotic associations in other subfamilies. For instance, Haematosiphoninae, characterized by broad morphological, ecological, and host diversity across the Americas, is represented here by only two *Sodalis*-harbouring species. Expanding sampling in this group could clarify the coevolutionary history of *Sodalis* and potentially uncover novel symbiotic associations.

Likewise, deeper insight into the relationship between Cacodminae and *Symbiopectobacterium* may be achieved by including additional taxa, such as *Aphrania* (a genus related to *Cacodmus*) and the species rich genera *Stricticimex* and *Loxaspis*, both related to *Leptocimex*. An especially intriguing direction is to extend analyses to the Polyctenidae, a group recently proposed to have evolved within Cimicidae (Szentivanyi et al. 2022). Polyctenids are more specialized than cimicids, spending their entire life cycle on bat host (an adaptation supported by vivipary). Comparing cimicids and polyctendis may shed light on how ecology and specialization influence the dynamics of symbiont acquisition and the stability of symbiont associations. Another important goal is to conduct population-level screenings to determine which of the detected strains are fixed in their host species. Given the primary nutritional role of the symbionts, it is especially important to identify which compounds are truly essential for cimicid hosts, particularly among B vitamins. In this study, some consistent patterns emerged, such as the universal production of riboflavin and the widespread degeneration of the thiamine biosynthesis pathway. A similar pattern has been reported across several *Arsenophonus* strains associated with obligate blood-feeding flies in the family Hippoboscidae (Martin Říhová et al. 2023). On the other hand, inconsistencies in other vitamins are notable and warrants further investigation. A particularly illustrative case is biotin. Multiple lines of evidence suggest that biotin synthesis is essential for symbiont function (e.g., the repeated acquisition of the biosynthetic pathway via horizontal transfer in symbiotic bacteria, or the pathway’s duplication observed in this study). However, this is difficult to reconcile with the apparent nonfunctionality of the biotin pathway in some symbionts. This discrepancy may reflect limitations in our ability to confidently assess pathway functionality, or it may indicate different nutritional requirements among host species. Resolving these questions will likely require complementary transcriptomic and experimental approaches beyond genomic data alone.

## Methods

### Samples and DNA preparation

Specimens of cimicids were collected from various localities across four continents between 2005 and 2015 (Supplementary table S1). For 16S rRNA amplicon analysis, DNA was extracted from 35 individuals using the E.Z.N.A. Insect DNA Kit (Omega BIO-TEK). Based on amplicon profiles, DNA concentration, and integrity, 14 samples representing 13 different species (*C. lectularius* was represented by two samples, one from human host, the other from bat) were selected for metagenomic analysis (Supplementary table S4). DNA concentrations were measured using a Qubit 2.0 Fluorometer (Invitrogen, Carlsbad, CA, USA), and integrity was assessed by agarose gel electrophoresis. To enrich bacterial DNA, host methylated DNA was selectively removed using NEBNext® Microbiome DNA Enrichment Kit (New England BioLabs). Final DNA concentrations were again quantified using the Qubit 2.0 Fluorometer with High Sensitivity reagents.

### Amplicons

Using the QIAGEN Multiplex PCR Kit (Qiagen, Hilden, Germany), we amplified the V3–V4 region of the 16S rRNA gene from all samples, as well as from four blank (negative) PCR controls and two commercial gDNA (positive) controls (ATCC® MSA-1000™ and MSA-1001™, each containing the same ten bacterial species in different proportions). Two-step PCR was performed with 341F and 805R primers, containing staggered spacer and Illumina overhang adapters, followed by index PCR to add sample-specific barcodes (Illumina 16S Metagenomic Sequencing Library Preparation Guide). Purified amplicons were quantified, pooled equimolarly, and sequenced on an Illumina MiSeq using v2 chemistry (2 × 250 bp paired-end reads).

Raw Fastq files were processed using USEARCH v11.0.667 (Edgar 2013), including primer stripping, read merging, trimming, quality filtering, and OTU picking. Reads were merged allowing for zero mismatches within the alignment. The merged reads were quality filtered with a stringent option -fastq_maxee set to 0.5, and trimmed to 400 bp. OTU table was generated by clustering these sequences at 100% identity, followed by de novo OTU picking USEARCH global alignment at 97% identity, including chimera removal (Edgar 2013). Taxonomic assignment was conducted via BLASTn against the SILVA_138.2 database (https://www.arb-silva.de/no_cache/download/archive/release_138_2/Exports/). Data filtering, rarefaction and heat map visualization were carried out in R using microeco v0.16.0 (Liu et al. 2021) and ggplot2 v3.4.2 (Ito and Murphy 2013). In detail, OTU table was taxonomically filtered to remove archaeal, eukaryotic, mitochondrial, and chloroplast OTUs. We expected symbiont OTUs to be abundant, so any OTU comprising <1 % of reads in a sample was set to zero for that sample (presence across other samples was not considered). Finally, the dataset was rarefied to 1 000 reads per sample, and samples with fewer reads were excluded from further analysis.

### Metagenomics

The shotgun genomic libraries were prepared from enriched gDNAs, multiplexed and sequenced on NovaSeq 6000 from both ends. Metagenomic raw reads were quality trimmed using Trimmomatic (Bolger, Lohse and Usadel 2014) with -phred33 option and assembled with SPades (Bankevich et al. 2012) using the parameters --meta -t 50 -k 21,33,55,77,99,127. For several samples, the assembly failed due to memory allocation issues, likely caused by high complexity of the data. In these cases, the data were downsampled to 50% using seqtk (https://github.com/lh3/seqtk). From each successful assembly, a BLAST database was created and screened for contigs corresponding to symbionts indicated by the amplicon data: *Wolbachia*, *Symbiopectobacterium*, *Sodalis*, *Serratia,* and *Tisiphia*. To obtain representative queries, we downloaded genomes from NCBI that covered the diversity of each of these five bacterial taxa (Supplementary table S4), and used Roary program (Page et al. 2015) to construct pangenomes in which each gene was represented only once. These datasets were used as BLAST queries against each assembly-derived database. Contigs retrieved by these queries were annotated using PROKKA (Seemann 2014), and their taxonomic origins were validated using a “back-BLAST” procedure: all annotations from each contig were extracted and compared against NCBI nr database. For each contig, the bacterial genus with the highest number of hits was considered as the contig’s taxonomic origin; contigs with predominant eukaryotic hits were excluded from further analysis. This procedure yielded sets of contigs putatively representing five symbiotic genera: *Wolbachia*, *Sodalis*, *Symbiopectobacterium*, *Serratia,* and *Tisiphia*.

#### Double-infections

As predicted by the amplicon results, three samples (C51–*Cimex hirundinis*, C12– *Cyanolicimex patagonicus*, and C21–*Psitticimex uritui*) each harbored two distinct *Wolbachia* strains. Metagenomic data provided two additional lines of evidence. First, Orthofinder results revealed an excess of duplicated orthologs in these samples, suggesting the presence of more than one *Wolbachia* genome. Second, BLAST searches using annotated *Wolbachia* genes as queries against the complete metagenomic assemblies retrieved most genes in two copies, located on contigs with distinct coverage. Because the two *Wolbachia* genomes within a single host sample could not be fully separated (due to overlapping contigs coverage), we employed different strategies depending on the analysis. For phylogenetic analysis, we prioritized selecting genes that reliably represented the two distinct genomes over maximizing the total gene count. To achieve this, contigs were grouped into “high coverage” and “low coverage” sets, excluding those with intermediate coverage. These two sets were treated as separate genomes in a new Orthofinder run and subsequent phylogenetic analyses. For metabolic analyses, we used the complete set of all *Wolbachia* genes from a sample as a single “composite genome” to ensure inclusion of all genes potentially involved in the metabolic pathways reconstructed for the sample. The main genomic characteristics (Table 1) were derived mostly from the PROKKA annotations. Additionally, we used PHASTEST (Wishart et al. 2023) as an alternative tool to identify phage-associated sequences and Pseudofinder (Syberg-Olsen et al. 2022) to quantify pseudogenes.

### Phylogeny

Phylogenetic reconstructions were conducted for five sets: Cimicidae hosts, *Wolbachia*, *Symbiopectobacterium*, *Sodalis*, and *Serratia*. Due to the enrichment of bacterial sequences at the expense of host DNA (see above), we did not base the Cimicidae phylogeny on genome drafts. Instead, to obtain representative sets of orthologous sequences, we adopted the following strategy: coding sequences (CDSs) were extracted from the reference genome of *Cimex lectularius* (GCF_000648675.2), and used as queries to BLAST the assembly databases of the cimicid samples. Open reading frames (ORFs) in the retrieved contigs were identified using the built-in algorithm of Geneious prime (Kearse et al. 2012). These sequence sets were then processed using Orthofinder (Emms and Kelly 2019) to identify orthologous sequences. As an outgroup, we included the reduviid species *Rhynocoris fuscipes* (NCBI accession: GCA_040020575.1), for which ORFs were identified using the same method. Single-copy orthologs were aligned using MAFFT (Katoh and Standley 2013) with default settings. Alignments were concatenated, and unreliably aligned regions were removed using GBlocks (Talavera and Castresana 2007) with default parameters. Phylogenies were inferred using two approaches. First, Bayesian inference (BI) was performed in PhyloBayes MPI (Lartillot et al. 2013), running two independent chains under the CAT-GTR model, with the number of generations determined by convergence parameter (maxdiff < 0.1). Second, maximum likelihood (ML) analysis was carried out in IQtree (Trifinopoulos et al. 2016), with matrices partitioned by gene. The best partitioning scheme and substitution models were automatically, and node support was assessed using 1,000 ultrafast bootstrap replicates. For the four bacterial datasets, phylogenetic reconstructions were based on their annotated genome drafts. The entire workflow, from ortholog identification to alignments and phylogenetic analyses, followed the same procedure as described above for the cimicid hosts. The trees were visualized in the program iTOL (Letunic and Bork 2021). For *Tisiphia,* with only fragmentary low-coverage genome draft, we based the phylogenetic placement on the NCBI fast minimum evolution algorithm, using two different genes in independent analyses (Supplementary figure S4).

### Metabolic analysis

Metabolic capacities were assessed for 22 samples (Supplementary table S4). In addition to the newly assembled symbiont genomes, two reference strains, *Symbiopectobacterium purcellii* (NZ_CP081864) and *Sodalis praecaptivus* (GCF_039646275.1), were included for comparison. In samples with *Wolbachia* double infections (C51-*Cimex hirundinis,* C12-*Cyanolicimex patagonicus, and* C21-*Psitticimex uritui*), the two genomes were treated as a single entry by combining all *Wolbachia* genes into one set per host (see above). Metabolic functions were evaluated using the Kyoto Encyclopedia of Genes and Genomes (KEGG) database (Kanehisa et al. 2016a). K numbers linking genes to metabolic functions were assigned to all PROKKA-annotated CDSs by BlastKOALA server (Kanehisa, Sato and Morishima 2016b), and the resulting metabolic capacities were mapped to KEGG-defined metabolic pathways. Particular emphasis was placed on evaluating the functionality of B-vitamin biosynthetic pathways. Following the approach from our previous work (Martin Říhová et al. 2023), each missing gene was assessed individually. Two criteria were considered: (i) gene absent across all or most symbionts within otherwise intact pathways were assumed to be potentially non-essential; and (ii) pathways lacking a gene known to have alternative enzymes were also considered potentially functional.

## Data availability

Mendeley Data Reserved DOI: 10.17632/ny4ymjsxtd.1. Illumina reads and genome drafts were deposited in the NCBI; all accession numbers are available in Supplementary tables S1 and S4.

## Acknowledgments

Part of the computational resources were provided by the e-INFRA CZ project (ID:90254), supported by the Ministry of Education, Youth and Sports of the Czech Republic. This work was supported by the Grant Agency of the Czech Republic (grant number 24-10943S to V.H.).

## References

Alickovic, L., K. Johnson & B. Boyd (2021) The reduced genome of a heritable symbiont from an ectoparasitic feather feeding louse. Bmc Ecology and Evolution, 21, 108. 10.1186/s12862-021-01840-7.

Balvin, O., S. Roth, B. Talbot & K. Reinhardt (2018) Co-speciation in bedbug *Wolbachia* parallel the pattern in nematode hosts. Scientific Reports, 8, 8797. 10.1038/s41598-018-25545-y

Bankevich, A., S. Nurk, D. Antipov, A. A. Gurevich, M. Dvorkin, A. S. Kulikov, V. M. Lesin, S. I. Nikolenko, S. Pham, A. D. Prjibelski, A. V. Pyshkin, A. V. Sirotkin, N. Vyahhi, G. Tesler, M. A. Alekseyev & P. A. Pevzner (2012) SPAdes: a new genome assembly algorithm and its applications to single-cell sequencing. J Comput Biol, 19, 455–77.

Benoit, J. B., Z. N. Adelman, K. Reinhardt, A. Dolan, M. Poelchau, E. C. Jennings, E. M. Szuter, R. W. Hagan, H. Gujar, J. N. Shukla, F. Zhu, M. Mohan, D. R. Nelson, A. J. Rosendale, C. Derst, V. Resnik, S. Wernig, P. Menegazzi, C. Wegener, N. Peschel, J. M. Hendershot, W. Blenau, R. Predel, P. R. Johnston, P. Ioannidis, R. M. Waterhouse, R. Nauen, C. Schorn, M. C. Ott, F. Maiwald, J. S. Johnston, A. D. Gondhalekar, M. E. Scharf, B. F. Peterson, K. R. Raje, B. A. Hottel, D. Armisen, A. J. J. Crumiere, P. N. Refki, M. E. Santos, E. Sghaier, S. Viala, A. Khila, S. J. Ahn, C. Childers, C. Y. Lee, H. Lin, D. S. T. Hughes, E. J. Duncan, S. C. Murali, J. X. Qu, S. Dugan, S. L. Lee, H. Chao, H. Dinh, Y. Han, H. Doddapaneni, K. C. Worley, D. M. Muzny, D. Wheeler, K. A. Panfilio, I. M. V. Jentzsch, E. L. Vargo, W. Booth, M. Friedrich, M. T. Weirauch, M. A. E. Anderson, J. W. Jones, O. Mittapalli, C. Y. Zhao, J. J. Zhou, J. D. Evans, G. M. Attardo, H. M. Robertson, E. M. Zdobnov, J. M. C. Ribeiro, R. A. Gibbs, J. H. Werren, S. R. Palli, C. Schal & S. Richards (2016) Unique features of a global human ectoparasite identified through sequencing of the bed bug genome. Nature Communications, 7,10165. doi: 10.1038/ncomms10165. PMID: 26836814;

Bolger, A., M. Lohse & B. Usadel (2014) Trimmomatic: a flexible trimmer for Illumina sequence data. Bioinformatics, 30, 2114–2120.

Cagatay, N. S., M. Akhoundi, A. Izri, S. Brun & G. D. D. Hurst (2025) Prevalence of Heritable Symbionts in Parisian Bedbugs (Hemiptera: Cimicidae). Environmental Microbiology Reports, 17, e70054.

Douglas, A. E. 2015. Multiorganismal Insects: Diversity and Function of Resident Microorganisms. In Annual Review of Entomology*, Vol* 60, ed. M. R. Berenbaum, 17–34.

Duron, O. & Y. Gottlieb (2020) Convergence of Nutritional Symbioses in Obligate Blood Feeders. Trends in Parasitology, 36, 816–825.

Edgar, R. C. (2013) UPARSE: highly accurate OTU sequences from microbial amplicon reads. Nature Methods, 10, 996–8. doi: 10.1038/nmeth.2604

Emms, D. & S. Kelly (2019) OrthoFinder: phylogenetic orthology inference for comparative genomics. Genome Biology, 20. DOI: 10.1186/s13059-019-1832-y.

Hickin, M., M. Kakumanu & C. Schal (2022) Effects of *Wolbachia* elimination and B-vitamin supplementation on bed bug development and reproduction. Scientific Reports, 12, 10270. 10.1038/s41598-022-14505-2

Hosokawa, T., R. Koga, Y. Kikuchi, X. Y. Meng & T. Fukatsu (2010) Wolbachia as a bacteriocyte-associated nutritional mutualist. Proceedings of the National Academy of Sciences of the United States of America, 107, 769–774.

Husnik, F. & J. P. McCutcheon (2016) Repeated replacement of an intrabacterial symbiont in the tripartite nested mealybug symbiosis. Proceedings of the National Academy of Sciences of the United States of America, 113, E5416–E5424.

Hypsa, V. & S. Aksoy (1997) Phylogenetic characterization of two transovarially transmitted endosymbionts of the bedbug Cimex lectularius (Heteroptera: Cimicidae). Insect Molecular Biology, 6, 301–304.

Ito, K. & D. Murphy (2013) Application of ggplot2 to Pharmacometric Graphics. CPT Pharmacometrics Syst Pharmacol, 2, e79.

Kanehisa, M., Y. Sato, M. Kawashima, M. Furumichi & M. Tanabe (2016a) KEGG as a reference resource for gene and protein annotation. Nucleic Acids Research, 44, D457–D462.

Kanehisa, M., Y. Sato & K. Morishima (2016b) BlastKOALA and GhostKOALA: KEGG Tools for Functional Characterization of Genome and Metagenome Sequences. Journal of Molecular Biology, 428, 726–731.

Katoh, K. & D. Standley (2013) MAFFT Multiple Sequence Alignment Software Version 7: Improvements in Performance and Usability. Molecular Biology and Evolution, 30, 772–780.

Kearse, M., R. Moir, A. Wilson, S. Stones-Havas, M. Cheung, S. Sturrock, S. Buxton, A. Cooper, S. Markowitz, C. Duran, T. Thierer, B. Ashton, P. Meintjes & A. Drummond (2012) Geneious Basic: An integrated and extendable desktop software platform for the organization and analysis of sequence data. Bioinformatics, 28, 1647–1649.

Lartillot, N., N. Rodrigue, D. Stubbs & J. Richer (2013) PhyloBayes MPI: Phylogenetic Reconstruction with Infinite Mixtures of Profiles in a Parallel Environment. Systematic Biology, 62, 611–615.

Letunic, I. & P. Bork (2021) Interactive Tree Of Life (iTOL) v5: an online tool for phylogenetic tree display and annotation. Nucleic Acids Research, 49, W293–W296.

Liu, C., Y. Cui, X. Li & M. Yao (2021) microeco: an R package for data mining in microbial community ecology. Fems Microbiology Ecology, 97, fiaa255., 10.1093/femsec/fiaa255

Martin Říhová, J., S. Gupta, A. Darby, E. Nováková & V. Hypša (2023) *Arsenophonus* symbiosis with louse flies: multiple origins, coevolutionary dynamics, and metabolic significance. mSystems, 0, e00706–23.

Moran, N. & P. Baumann (2000) Bacterial endosymbionts in animals. Current Opinion in Microbiology, 3, 270–275.

Moriyama, M., N. Nikoh, T. Hosokawa & T. Fukatsu (2015) Riboflavin Provisioning Underlies Wolbachia’s Fitness Contribution to Its Insect Host. Mbio, 6, e01732–15. doi: 10.1128/mBio.01732-15. PMID: 26556278; PMCID: PMC4659472.

Nadal-Jimenez, P., S. Siozios, N. Halliday, M. Camara & G. Hurst (2022) *Symbiopectobacterium purcellii*, gen. nov., sp. nov., isolated from the leafhopper Empoasca decipiens. International Journal of Systematic and Evolutionary Microbiology, 72. doi: 10.1099/ijsem.0.005440. PMID: 35695864.

Nikoh, N., T. Hosokawa, M. Moriyama, K. Oshima, M. Hattori & T. Fukatsu (2014) Evolutionary origin of insect-Wolbachia nutritional mutualism. Proceedings of the National Academy of Sciences of the United States of America, 111, 10257–10262.

Oakeson, K. F., R. Gil, A. L. Clayton, D. M. Dunn, A. C. von Niederhausern, C. Hamil, A. Aoyagi, B. Duval, A. Baca, F. J. Silva, A. Vallier, D. G. Jackson, A. Latorre, R. B. Weiss, A. Heddi, A. Moya & C. Dale (2014) Genome Degeneration and Adaptation in a Nascent Stage of Symbiosis. Genome Biology and Evolution, 6, 76–93.

Page, A. J., C. A. Cummins, M. Hunt, V. K. Wong, S. Reuter, M. T. Holden, M. Fookes, D. Falush, J. A. Keane & J. Parkhill (2015) Roary: rapid large-scale prokaryote pan genome analysis. Bioinformatics, 31, 3691–3.

Poulain, M., E. Rosinski, H. Henri, S. Balmand, M. Delignette-Muller, A. Heddi, R. Lasseur, F. Vavre, A. Zaidman-Rémy & N. Kremer (2024) Development, feeding, and sex shape the relative quantity of the nutritional obligatory symbiont *Wolbachia* in bed bugs. Frontiers in Microbiology, 15. 10.3389/fmicb.2024.1386458

Rasgon, J. & T. Scott (2004) Phylogenetic characterization of *Wolbachia* symbionts infecting *Cimex-lectularius* L. and *Oeciacus vicarius* Horvath (Hemiptera: Cimicidae). Journal of Medical Entomology, 41, 1175–1178.

Reinhardt, K. & M. Siva-Jothy (2007) Biology of the bed bugs (Cimicidae). Annual Review of Entomology, 52, 351–374.

Rihova, J., E. Novakova, F. Husnik & V. Hypsa (2017) *Legionella* Becoming a Mutualist: Adaptive Processes Shaping the Genome of Symbiont in the Louse Polyplax serrata. Genome Biology and Evolution, 9, 2946–2957.

Rihova, J., R. Vodicka & V. Hypsa (2025) An obligate symbiont of *Haematomyzus elephantis* with a strongly reduced genome resembles symbiotic bacteria in sucking lice. Applied and Environmental Microbiology. 0:e00220–25. 10.1128/aem.00220-25

Rio, R., G. Attardo & B. Weiss (2016) Grandeur Alliances: Symbiont Metabolic Integration and Oblicate Arthopod Hematophagy. Trends in Parasitology, 32, 739–749.

Roth, S., O. Balvin, M. Siva-Jothy, O. Di Iorio, P. Benda, O. Calva, E. Faundez, F. Khan, M. McFadzen, M. Lehnert, R. Naylor, N. Simov, E. Morrow, E. Willassen & K. Reinhardt (2019) Bedbugs Evolved before Their Bat Hosts and Did Not Co-speciate with Ancient Humans. Current Biology, 29, 1847–1853.e4. doi: 10.1016/j.cub.2019.04.048.

Ríhová, J., G. Batani, S. Rodríguez-Ruano, J. Martinu, F. Vácha, E. Nováková & V. Hypsa (2021) A new symbiotic lineage related to *Neisseria* and *Snodgrassella* arises from the dynamic and diverse microbiomes in sucking lice. Molecular Ecology, 30, 2178–2196.

Sakamoto, J., J. Feinstein & J. Rasgon (2006) *Wolbachia* infections in the Cimicidae:: Museum specimens as an untapped resource for endosymbiont surveys. Applied and Environmental Microbiology, 72, 3161–3167.

Seemann, T. (2014) Prokka: rapid prokaryotic genome annotation. Bioinformatics, 30, 2068–2069.

Sochova, E., F. Husnik, E. Novakova, A. Halajian & V. Hypsa (2017) *Arsenophonus* and *Sodalis* replacements shape evolution of symbiosis in louse flies. Peerj, 5, e4099. doi: 10.7717/peerj.4099.

Sudakaran, S., C. Kost & M. Kaltenpoth (2017) Symbiont Acquisition and Replacement as a Source of Ecological Innovation. Trends in Microbiology, 25, 375–390.

Syberg-Olsen, M. J., A. I. Garber, P. J. Keeling, J. P. McCutcheon & F. Husnik (2022) Pseudofinder: detection of pseudogenes in prokaryotic genomes. Molecular Biology and Evolution. 39, msac153. 10.1093/molbev/msac153

Szentivanyi, T., S. Hornok, A. Kovacs, N. Takacs, M. Gyuranecz, W. Markotter, P. Christe & O. Glaizot (2022) Polyctenidae (Hemiptera: Cimicoidea) species in the Afrotropical region: Distribution, host specificity, and first insights to their molecular phylogeny. Ecology and Evolution, 12, e9357. doi: 10.1002/ece3.9357.

Talavera, G. & J. Castresana (2007) Improvement of phylogenies after removing divergent and ambiguously aligned blocks from protein sequence alignments. Systematic Biology, 56, 564–577.

Thongprem, P., S. Evison, G. Hurst & O. Otti (2020) Transmission, Tropism, and Biological Impacts of Torix *Rickettsia* in the Common Bed Bug *Cimex lectularius* (Hemiptera: Cimicidae). Frontiers in Microbiology, 11, 608763. doi: 10.3389/fmicb.2020.608763.

Trifinopoulos, J., L. Nguyen, A. von Haeseler & B. Minh (2016) W-IQ-TREE: a fast online phylogenetic tool for maximum likelihood analysis. Nucleic Acids Research, 44, W232–W235.

Werren, J., L. Baldo & M. Clark (2008) *Wolbachia*: master manipulators of invertebrate biology. Nature Reviews Microbiology, 6, 741–751.

Wishart, D., S. Han, S. Saha, E. Oler, H. Peters, J. Grant, P. Stothard & V. Gautam (2023) PHASTEST: faster than PHASTER, better than PHAST. Nucleic Acids Research, 51, W443–W450.

