## Supplementary material for "Dynamic but constrained: Repeated acquisitions of nutritional symbionts in bed bugs (Heteroptera: Cimicidae) from a narrow taxonomic pool": Suplementary figures

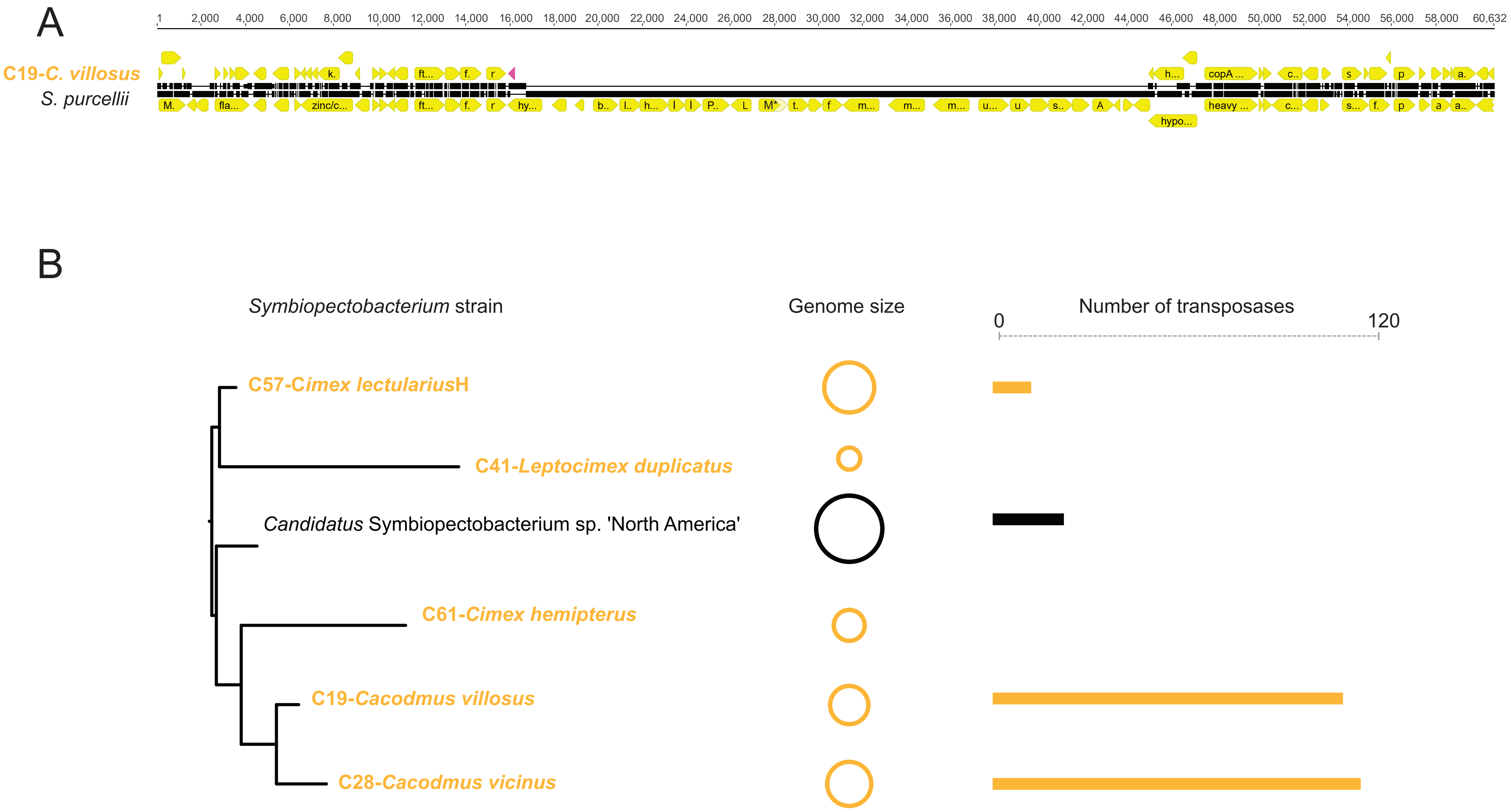


**Supplementary figure S1**. A: An example of gene loss in C19-*Cacodmus villosus* genome compared to *Symbiopectobacterium purcelii*. Yellow shapes=CDSs, pink triangle=transposase. B: Comparison of genome sizes and transposase numbers among the *Symbiopectobacterium* strains. Detailed genome characteristics are provided in Table 1.

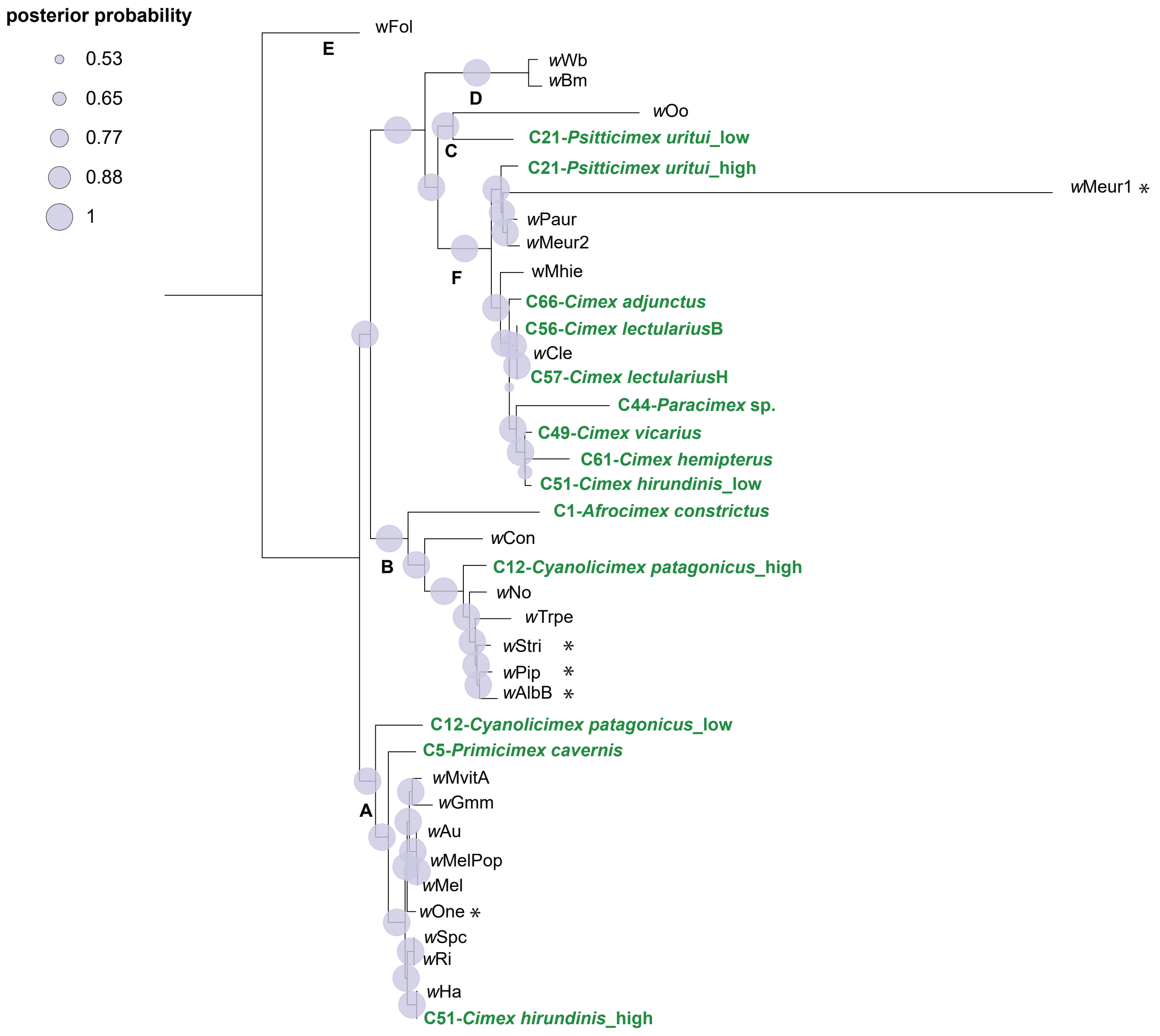


**Supplementary figure S2**. Phylogenetic relationships inferred by BI from concatenated matrix of 115 single-copy orthologs (27,850 amino acid residues). Letters at the nodes indicate *Wolbachia* supergroups. Strains identified in this study from cimicid species are highlighted in bold green. The designations "high" and "low" indicate differences in sequencing coverage between co-occurring strains in the same host. Asterisks mark branches whose positions differ in ML analysis. Accession numbers for all included taxa are provided in Supplementary Table SX.

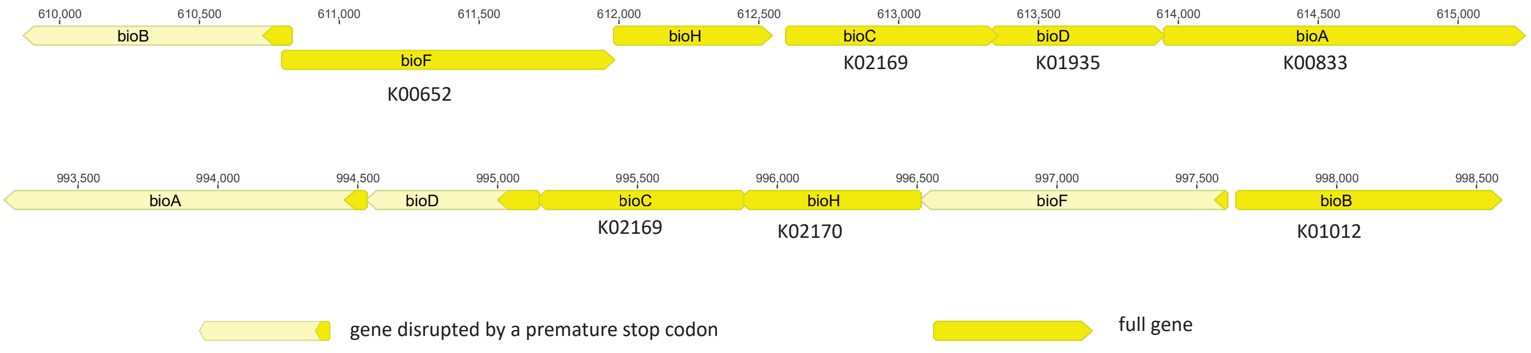


**Supplementary figure S3**. Two biotin operons in genome of C44-Paracimex with complementary disrupted/functional genes. The numbers above genes indicate positions in the genomes; the assigned K numbers indicate which genes were recognized by BlastKoala as functional.

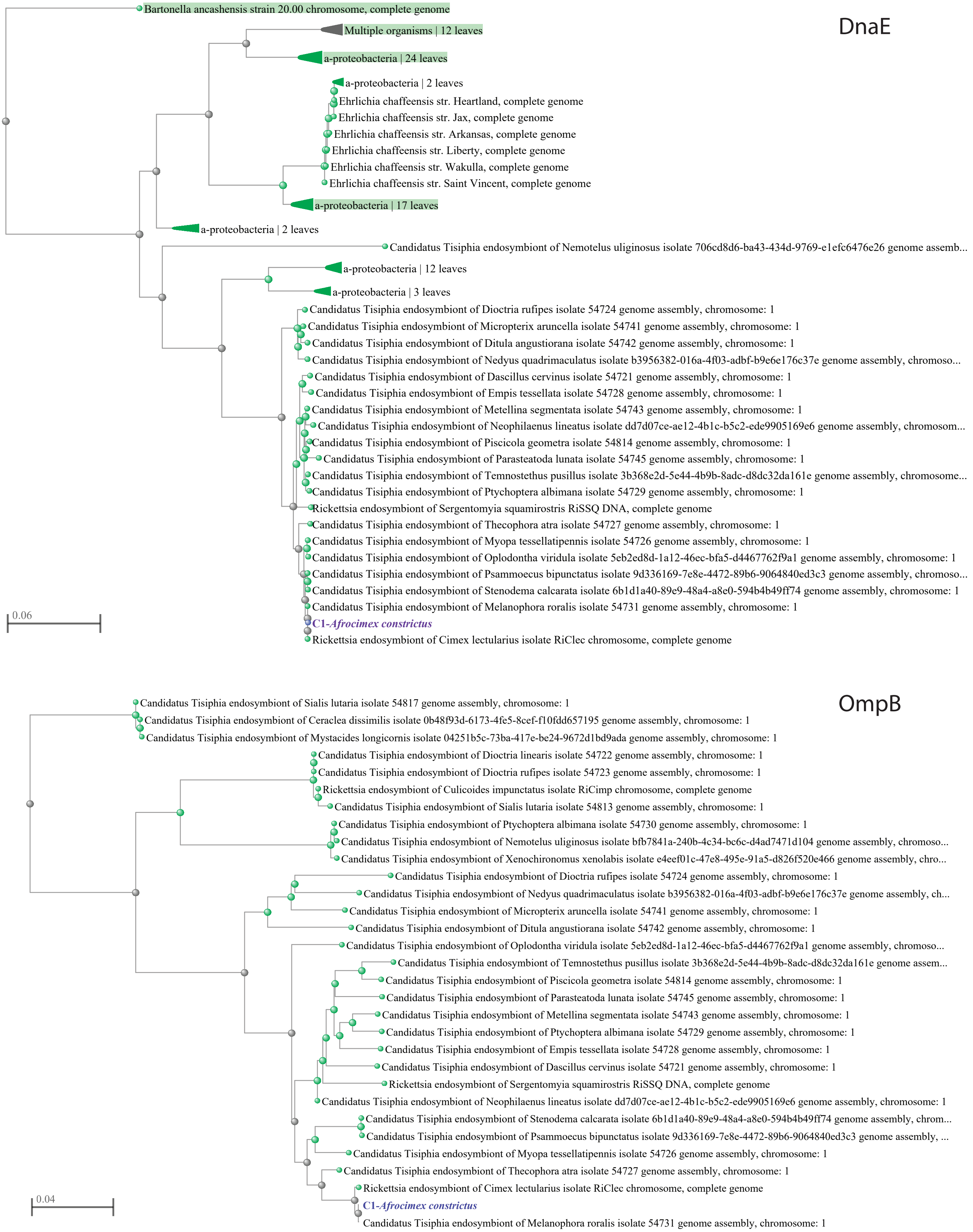


**Supplementary figure S4**. Phylogenetic trees inferred using NCBI-based fast minimum evolution algorithm, based on two genes (DnaE and OmpB) from fragments of the *Tisiphia* genome extracted from the C1-*Afrocimex constric-tus* metagenome.
